# Enhanced microbiota profiling in patients with quiescent Crohn’s disease through comparison with paired healthy first-degree relatives using fecal metagenomics and metabolomics

**DOI:** 10.1101/2024.02.24.581863

**Authors:** Wanning Chen, Yichen Li, Wenxia Wang, Sheng Gao, Dingfeng Wu, Na Jiao, Tao Xu, Min Zhi, Lixin Zhu, Ruixin Zhu

## Abstract

Prior studies indicate no correlation between gut microbiota of healthy first-degree relatives (HFDRs) of Crohn’s disease (CD) patients and development of CD. Here, we utilized HFDRs as controls to examine the microbiota and metabolome in individuals with active (CD-A) and quiescent (CD-R) CD, thereby minimizing the influence of genetic and environmental factors. Compared to non-relative controls, the use of HFDR controls identified fewer differential taxa. *Faecalibacterium, Dorea,* and *Fusicatenibacter* showed decreased abundances in CD-R, independent of inflammation, and correlated with fecal SCFAs. Validation with a large multi-center cohort confirmed decreased abundances in *Faecalibacterium* and other SCFA-producing genera in CD-R. Classification models based on these genera distinguished CD-R and CD-A from healthy individuals, in both the discovery and validation cohorts. Thus, the decreased presence of *Faecalibacterium, Dorea,* and *Fusicatenibacter* in CD-R likely contributed to disease relapse through reduced SCFA production, highlighting their potential as diagnostic markers and therapeutic targets for CD.

## Introduction

Crohn’s disease (CD), one of the major forms of inflammatory bowel diseases (IBD), is characterized by frequent relapses separated by periods of remission. The presence of gut microbiota is a fundamental requirement for CD development, as evidenced in studies involving both CD patients (Harper et al., 1985) and animal models(Sadlack et al., 1993; Sellon et al., 1998) of the illness.

A causal role of the gut microbiota in CD pathogenesis is supported by microbiota intervention studies with colitis patients(Gionchetti et al., 2000) and a mouse model of colities(Elinav et al., 2011). This is further supported by the presence of humoral(Arnott et al., 2004; Landers et al., 2002; Lee et al., 2021) and cellular(Morgan et al., 2022) immune responses against gut microbes in CD patients.

Given the key role of the gut microbiota in CD, coupled with the fact that patients with CD in remission (CD-R) frequently develop active colitis (CD-A)(Pauwels et al., 2022; Sutherland, 2000), it is reasonable to hypothesize that abnormal gut microbiota in CD-R might contribute more significantly to intestinal inflammation development than microbial features unique to CD-A. Therefore, many efforts have been spent to determine the gut microbial feature in CD-R. However, conflicting results were reported on gut microbial composition in CD-R. Between CD-R and healthy controls, differential taxa were observed by Galazzo et al.(Galazzo et al., 2019) and Pascal et al.(Pascal et al., 2017), but not by Sokol et al. (Sokol et al., 2009). Between CD-R and CD-A, different microbial composition and diversity were observed by Will et al.(Wills et al., 2014), but not by Pascal et al. (Pascal *et al*., 2017) or Halfvarson et al.(Halfvarson et al., 2017). Furthermore, for studies reporting differential taxa between CD-R and healthy controls, distinct observations were reported by different research groups (Galazzo *et al*., 2019; Pascal *et al*., 2017).The inconsistencies regarding the microbiota of CD-R might arise from the large impact of host genetic polymorphism(Cheng et al., 2021), environmental factors such as diet(Wu et al., 2011) and hygiene habits(Singhal et al., 2011). Increasing the sample size may not be a feasible solution to address confounding factors, as evidenced by previous studies with over 800 samples that reported divergent findings on microbial compositions in IBD(Imhann et al., 2018; Lloyd-Price et al., 2019; Pascal *et al*., 2017).

Inspired by a recent report that the gut microbial compositions of the healthy relatives of CD patients were not correlated with the development of CD(Raygoza Garay et al., 2023), we performed a microbiome and metabolome study on CD patients, including CD-A and CD-R, alongside paired healthy first-degree relative (HFDR) controls, so to minimize the influence of genetic and environmental confounding factors. In comparison to non-relative controls, the use of HFDR controls resulted in significantly reduced number of differential taxa in the microbiome of CD patients. Moreover, the gut dysbiosis in CD-R was not correlated with intestinal inflammation, and was characterized by a reduced ability to produce short-chain fatty acids (SCFAs). Additionally, the CD-R specific genera were proved to be valuable in distinguishing CD-R and CD-A from healthy controls. These observations were validated using a whole-genome sequencing dataset from a large independent multi-center cohort.

## Results

### Impact of environmental and host genetic variations on gut microbiota composition in CD patients

The current study was based on the DamnIBD cohort (detailed in STAR Methods), in which every IBD patient was paired with a healthy first-degree relative (HFDR). A total of 52 pairs of CD patients with healthy controls were enrolled, including 27 CD patients with active inflammation and their paired siblings (CD-A/Sibling-A) and 25 CD patients in remission and their paired siblings (CD-R/Sibling-R). Patients and their paired HFDRs had been living together for a substantial period, and approximately half of the patients are still living together with their HFDRs. Additionally, an independent control group was enrolled, consisting 44 healthy subjects not related to the CD patients (Non-relative). Patients in the study groups (CD-A, Sibling-A, CD-R, Sibling-R, Non-relative) had similar ages, sex ratios and body mass indices (Supplementary Table S1). The average dietary intake of macro- and micronutrients was similar between the CD-R patients and their HFDRs, as well as between the CD-R patients and non-relative controls, determined by 24-hour dietary recall (Supplementary Table S2). On the other hand, the dietary intake of fiber, calcium, potassium, vitamin C, and bete-carotene was lower in CD-A patients compared to their HFDRs (Supplementary Table S2), suggesting that different dietary habits could be a significant confounding factor in the study of the microbiota of CD-A.

From the 148 fecal microbiome samples, 86,805 reads of 16S rRNA gene were sequenced and assigned to 13,616 Amplicon Sequence Variants (ASVs). After excluding rare ASVs found in only two or fewer samples, 2,492 ASVs remained and were classified into 223 genera across 89 families and 11 phyla (Figure 1A, Supplementary Figure S1A, B). Similar patterns of the microbial compositions were observed between the CD-A and the CD-R groups, as well as among the healthy control groups (Sibling-A, Sibling-R, and non-relative).

**Figure 1.**
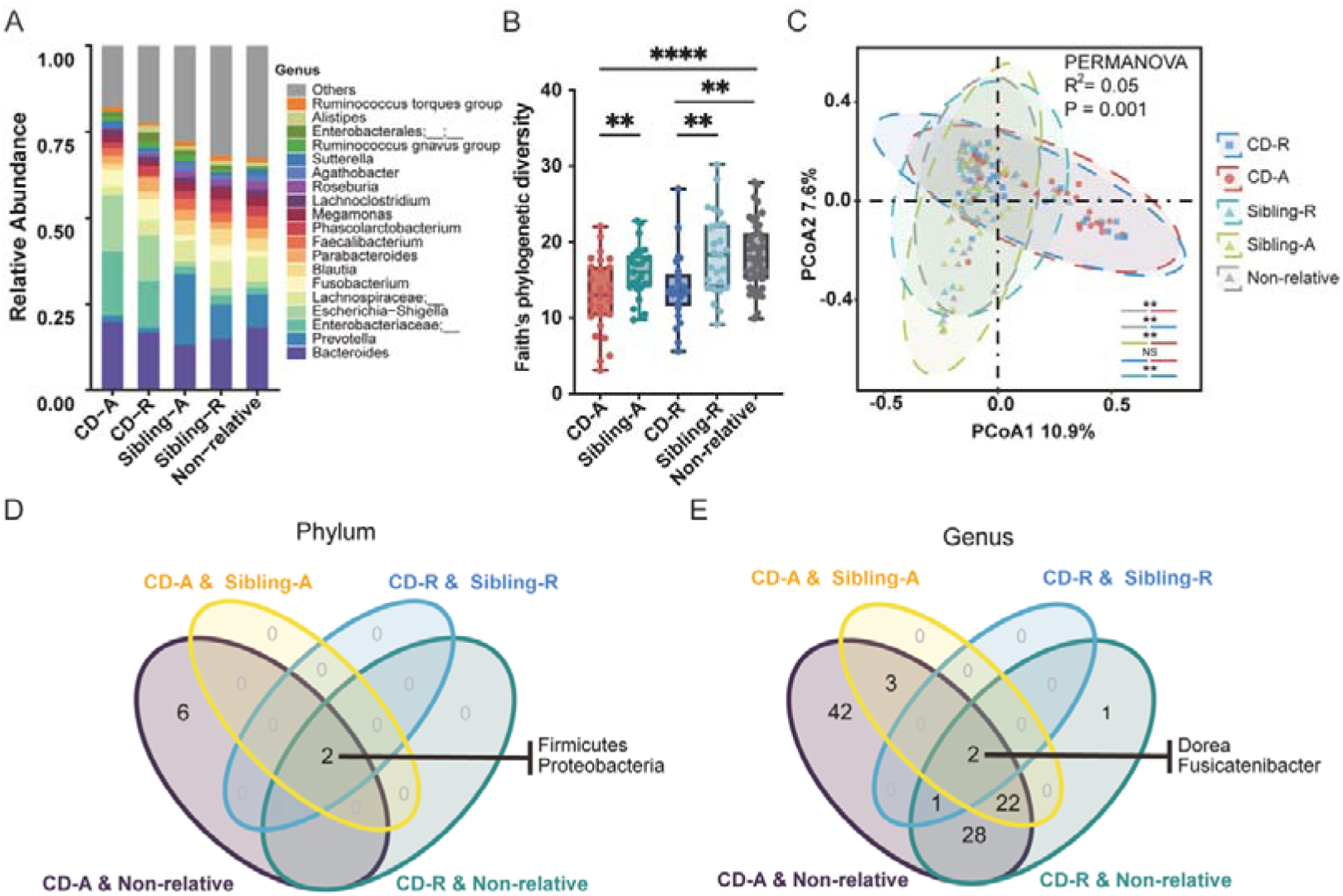
Microbial profiles in the gut of CD patients, their paired HFDRs, and non-relative controls: impact of environmental and host genetic variations on gut microbiota composition in CD patients. **(A)** Microbial compositions at the genus level across the study groups: patients with active Crohn’s disease (CD-A), patients with quiescent CD (CD-R), paired healthy siblings of CD-A patients (Sibling-A), paired healthy siblings of CD-R patients (Sibling-R), and non-relative healthy volunteers (non-relative). **(B)** Alpha diversities measured by Faith’s Phylogenetic Diversity (PD) in the gut of the study groups. Paired Wilcoxon rank sum tests were performed for CD-A vs. Sibling-A and CD-R vs. Sibling-R, and unpaired Wilcoxon rank sum tests for CD-A vs. non-relative and CD-R vs. non-relative. P values were adjusted for multiple tests. **, FDR < 0.01; ****: FDR < 0.0001. **(C)** Beta diversities assessed through PCoA analysis based on Bray-Curtis distance to evaluate compositional differences among the study groups. PERMANOVA was performed with P values adjusted for multiple tests. **, FDR < 0.01. **(D)** Venn diagram of the differential phyla, between CD-A and Sibling-A, CD-R and Sibling-R, CD-A and non-relative, as well as CD-R and non-relative. **(E)** Venn diagram of the differential genera.

The alpha and beta diversities of the study groups were assessed at the ASV level. When evaluating ecological diversities within each sample using a phylogenetic distance metric, both CD-A and CD-R groups exhibited lower diversities compared to their respective paired controls, while no significant difference was observed between the CD-A and the CD-R groups (Figure 1B). Similar results were obtained using Shannon index and other alpha-diversity metrics (Supplementary Figure S1B, C). Ecological diversities within the study groups were measured by UniFrac-based PCoA analysis, showing that the beta diversities of the CD-A and the CD-R groups were substantially different from those of the Sibling-A and the Sibling-R groups, respectively (Figure 1C). No notable divergence was observed between the CD-A and CD-R groups, nor between the Sibling-A and the Sibling-R groups.

Compositional changes were observed in the fecal microbiota at every taxonomic level in both the CD-A and the CD-R groups. At the phylum level, 8 and 2 differential phyla were observed in the CD-A and the CD-R groups, respectively, in comparison to the non-relative controls (Figure 1D). However, only Firmicutes and Proteobacteria were identified as differential phyla for both the CD-A and CD-R groups, when compared to their paired HFDR control groups (Sibling-A and Sibling-R) (Figure 1D). At the genus level, 98 and 54 differential genera were observed in the CD-A and CD-R groups, respectively, in comparison to the non-relative controls (Figure 1E, Supplementary Figure S2). Strikingly, when comparing with their paired HFDRs, the numbers of differential genera in the CD-A and CD-R groups were drastically reduced to 27 and 3, respectively (Figure 1E, Supplementary Figure S2). Similar observations were made at the family level (Supplementary Figure S2).

Thus, according to the numbers of differential taxa, the differences of the microbiota between CD patients and their HFDRs were smaller compared to the differences between CD patients and non-relative controls. To further evaluate the microbial distances of the study groups at the community level, UniFrac distances of the microbiota were assessed. Consistent with the results of differential taxa, the UniFrac distances between the CD and the paired HFDR groups were smaller than those between the CD and non-relative groups, for both active and quiescent CD (Supplementary Figure S3).

### Altered gut microbiota in patients with active and quiescent Crohn’s disease

The above results underscore the substantial impact of environmental and host genetic variations on the gut microbiota. In light of these results, paired HFDRs were employed as control subjects for further analyses of the microbiota in CD patients.

At the phylum level, Firmicutes, Bacteroidota, Proteobacteria and Fusobacteriota were the dominant phyla across all study groups. Both the CD-A and CD-R groups exhibited decreased abundances in Firmicutes, along with increased abundances in Proteobacteria, when compared to their paired HFDRs (Figure 2A).

**Figure 2.**
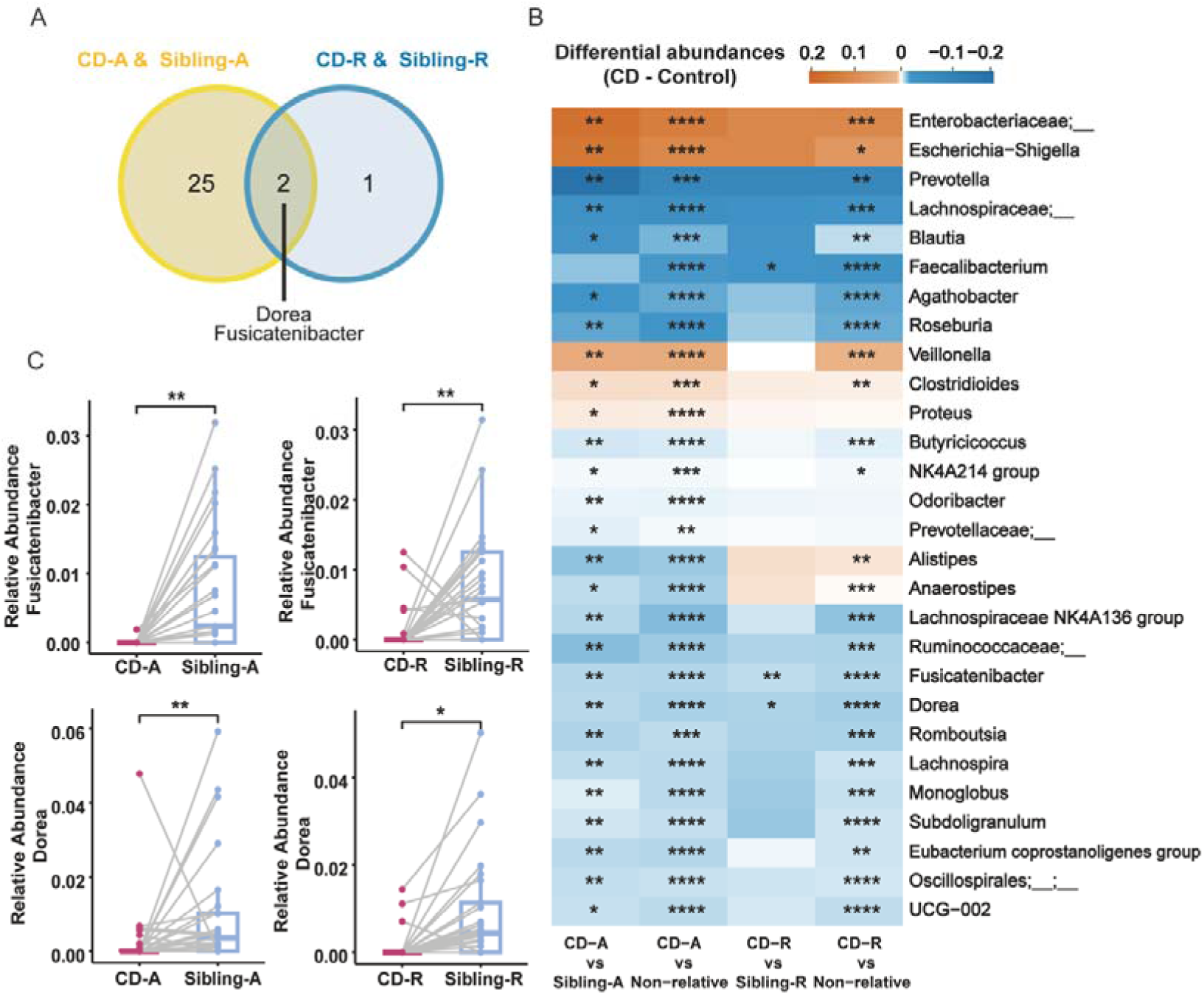
Differential genera in the gut of patients with active and quiescent CD. **(A)** Venn diagram of the differential genera between CD-A and Sibling-A, as well as CD-R and Sibling-R. The overlapping differential genera were indicated. **(B)** Heatmap displaying the differential abundances (DA) of genera between CD patients and healthy controls (CD - control). The heatmap encompasses four comparisons: CD-A vs. Sibling-A, CD-A vs. non-relative, CD-R vs. Sibling-R, and CD-R vs. non-relative. It includes all the differential genera identified in these comparisons. P values were adjusted for multiple tests. *, FDR < 0.1; **, FDR < 0.05; ***, FDR < 0.01; ****, FDR < 0.001. **(C)** Distribution of the relative abundances of the shared differential genera (*Dorea* and *Fusicatenibacter*) in CD and HFDR groups. Paired Wilcoxon rank sum tests were performed with P values adjusted for multiple tests. *, FDR < 0.1; **, FDR < 0.05.

At the genus level, 27 differential genera were found between the CD-A and the Sibling-A groups, whereas only 3 differential genera (*Dorea*, *Fusicatenibacter* and *Faecalibacterium*) were identified between the CD-R and the Sibling-R groups (Figure 2B, C). Among these differential genera, 25 were specific to CD-A, including *Escherichia-Shigella* (increased), an undefined genus of *Enterobacteriaceae* (increased), *Prevotella* (decreased) and *Roseburia* (decreased). *Faecalibacterium* (decreased) was the sole differential genus specific to CD-R. *Dorea* (decreased) and *Fusicatenibacter* (decreased) were common differential genera observed in both the CD-A and CD-R groups (Supplementary Table S3-4). According to the current hypothesis, the decreased abundances of *Dorea*, *Fusicatenibacter* and *Faecalibacterium* in CD-R may potentially contribute to future relapse of intestinal inflammation. On the other hand, the decreased abundances of *Dorea* and *Fusicatenibacter* in CD-A indicate their possible involvement in sustaining intestinal inflammation.

A notable pattern of gradually altered abundances from healthy controls to CD-R and then CD-A was observed for the majority of the differential genera in the CD-A group, indicating associations of these genera with the inflammatory activities (Figure 3). However, this pattern was not observed for two of the three differential genera (*Dorea* and *Faecalibacterium*) in CD-R (Figure 3), indicating that the alterations in these genera were not a consequence of inflammation.

**Figure 3.**
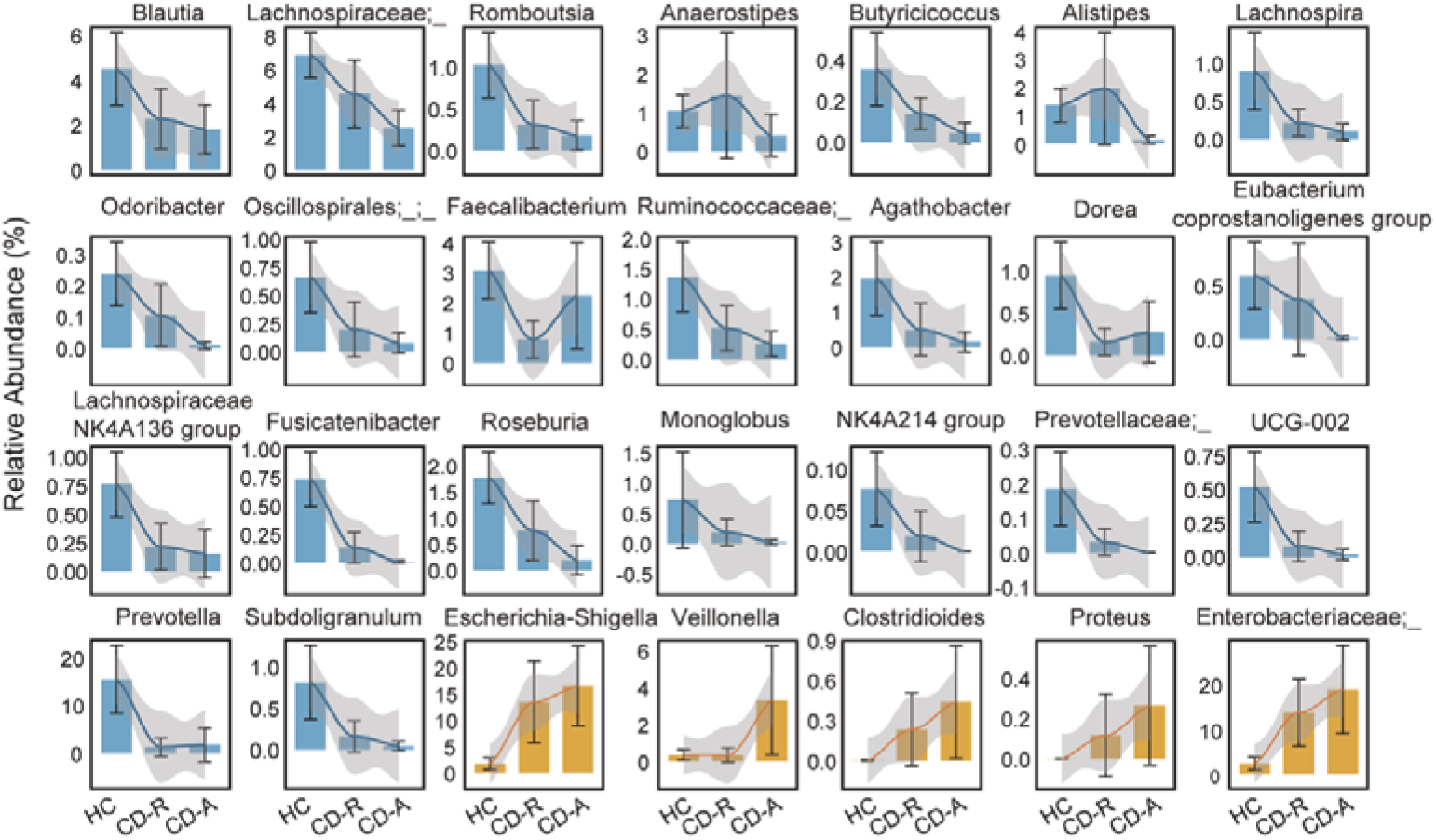
Comparison of the relative abundances of the differential genera among healthy controls, CD-R and CD-A groups. Plotted are average relative abundances of 28 differential genera in healthy controls, CD-R and CD-A groups. The healthy control group is the combination of the Sibling-A and Sibling-R groups. The error bars represent the 95% confidence interval. A Loess regression was applied to model the changing trends in relative abundances among the study groups. Blue bars indicate genera with decreased abundances in CD-A, while yellow bars increased abundances in CD-A.

To gain further insight into the relationship between microbial composition and the intestinal inflammation status, we examined possible associations between microbial genera and serum markers of inflammation, focusing on the 28 differential genera in CD-A and CD-R. The presence of significant correlations between microbial genera and serum inflammation markers was more frequently observed in CD-A compared to CD-R (Figure 4A). In CD-A, the differential genera *Escherichia-Shigella* correlated with four of the serum inflammation markers including hyper-sensitive C-reactive protein (hsCRP), CRP, platelet count, and erythrocyte sedimentation rate, followed by *Veillonella* correlating with three of the inflammation markers. In CD-R, no correlation was observed between any differential genera (*Dorea*, *Faecalibacterium* and *Fusicatenibacter*) and the serum inflammation markers.

**Figure 4.**
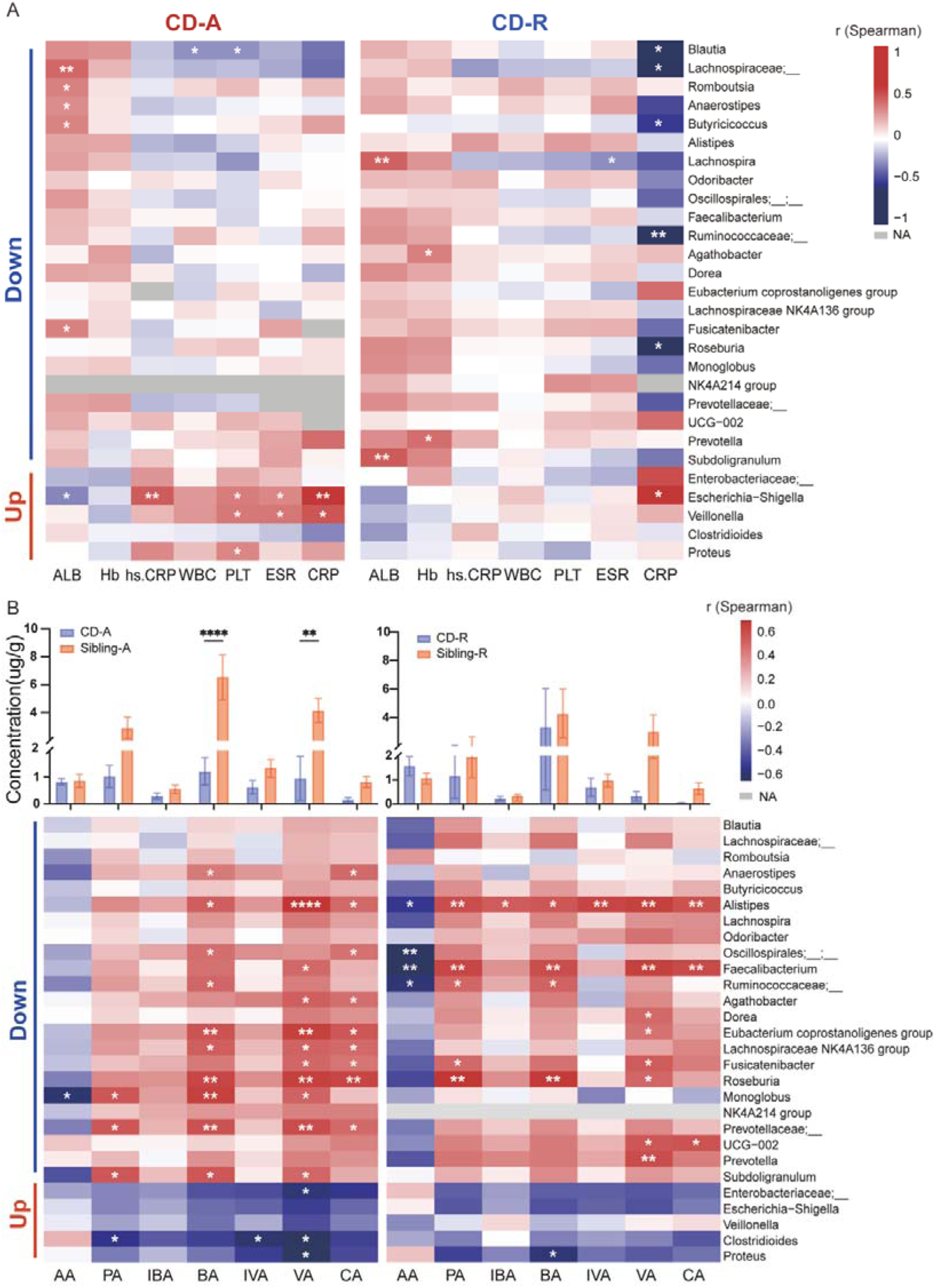
Correlations between microbial genera with serum inflammation markers and fecal short-chain fatty acids (SCFAs) **(A)** Correlations between microbial genera with serum inflammation markers in patients with CD. The heatmap represents the correlations between the relative abundances of differential genera and concentrations of serum inflammation markers. The Spearman coefficients (r) are color coded, and statistical significance indicated by asterisks. *, p < 0.05; **, p < 0.01. On the left side of the graph, differential genera were indicated for down- or up-regulated abundances in CD-A. Note that the CD-R specific differential genera (*Faecalibacterium, Dorea*, and *Fusicatenibacter*) did not show any correlation with the inflammation markers. ALB, albumin; Hb, hemoglobin; hs.CRP, hyper-sensitive C-reactive protein; WBC, while blood cell count; PLT, platelet count; ESR, erythrocyte sedimentation rate; CRP, C-reactive protein; NA, not applicable for correlation analysis. **(B)** Correlations between microbial genera with fecal SCFAs. The upper panels are fecal concentrations of SCFAs in the CD-A and CD-R groups, in comparison to their paired HFDRs (Sibling-A and Sibling-R groups), respectively. **, FDR < 0.01; ****, FDR < 0.0001. The lower panels are heatmaps of the Spearman correlations between the relative abundances of the 28 differential genera and SCFA concentrations, with the correlation coefficients color coded and p values indicated on the heatmap blocks. Data from both patients and their paired HFDRs are included in the analysis. *, p < 0.05; **, p < 0.01; ****, p < 0.0001.

These results have significant implications for understanding the causal relationship between the microbiota and inflammation in CD.

### Altered microbial functions in the CD-A and the CD-R groups

Functional analyses based on the 16S rRNA sequences revealed that the fecal microbiota of the CD-A were enriched in dozens of oxygen dependent metabolic pathways, including fatty acid beta-oxidation, sugar alcohol fermentation, and the TCA cycle, in comparison to their paired HFDR controls (Supplementary Figure S4). This enrichment correlated with higher abundances in facultative anaerobes, such as *Escherichia-Shigella*. Furthermore, elevated abundances in lipopolysaccharide biosynthesis in CD-A (Supplementary Figure S4) were consistent with an elevated representation of Gram-negative bacteria, such as Proteobacteria. Conversely, CD-A patients displayed decreased abundances in methanogenesis, a function typically associated with anaerobic bacteria (Supplementary Figure S4). This finding aligns with the observed decrease in anaerobic bacteria within CD-A group.

In contrast, much less differential pathways were identified in CD-R, which echoed the fact that less differential taxa were identified in CD-R compared to CD-A. Importantly, one of the top differential pathways, “pyruvate fermentation to acetate and lactate II”, was decreased in CD-R, as well as in CD-A (Supplementary Figure S4, Supplementary Table S5-6). Two other pathways involved in acetate production, “bifidobacterium shunt” and “heterolactic fermentation” were decreased in the CD-A group, and showed a trend of down-regulation in the CD-R group (Supplementary Figure S4). These data collectively suggest a reduced capability for acetate production within the gut of both the CD-R and CD-A groups.

### Correlations between fecal short-chain fatty acid (SCFA) levels and gut microbes in the CD-A and CD-R groups

The above results strongly indicate the relevance of SCFAs in the pathogenesis of CD. We thus examined the fecal levels of the SCFAs by targeted metabolomics analysis. CD-A group exhibited lower levels of butyrate and valerate compared to their paired HFDRs. Additionally, a trend of lower levels was observed in other SCFAs including acetate, propionate, iso-butyrate, iso-valerate and caproate (Figure 4B, upper left panel). Similarly, the CD-R group also exhibited trends of lower SCFA levels, including propionate, iso-butyrate, butyrate, iso-valerate, valerate, and caproate (Figure 4B, upper right panel).

Next, we performed correlation analyses and found associations between the abundances of several differential genera and SCFA levels. As expected, the CD-R specific differential genera, *Faecalibacterium* and *Fusicatenibacter*, showed correlations with multiple SCFAs (Figure 4B, lower panels). However, it was unexpected that *Butyricicoccus*, known for its butyrate production, did not correlate with butyrate levels, possibly due to the prevalence of other butyrate producing bacteria like *Faecalibacterium* in the gut(Louis and Flint, 2017). Interestingly, some of the differential genera (e.g. *Faecalibacterium*) were negatively correlated with acetate levels, likely due to the consumption of acetate by these bacteria.

### Potential diagnostic value of the differential genera

We next evaluated the diagnostic potential of CD-R specific microbial markers, namely *Dorea*, *Fusicatenibacter* and *Faecalibacterium*. A classification model was constructed with these three genera and achieved an AUROC of 0.82 for distinguishing CD-R from Sibling-R, and 0.88 for distinguishing CD-R from non-relative controls (Figure 5).

**Figure 5.**
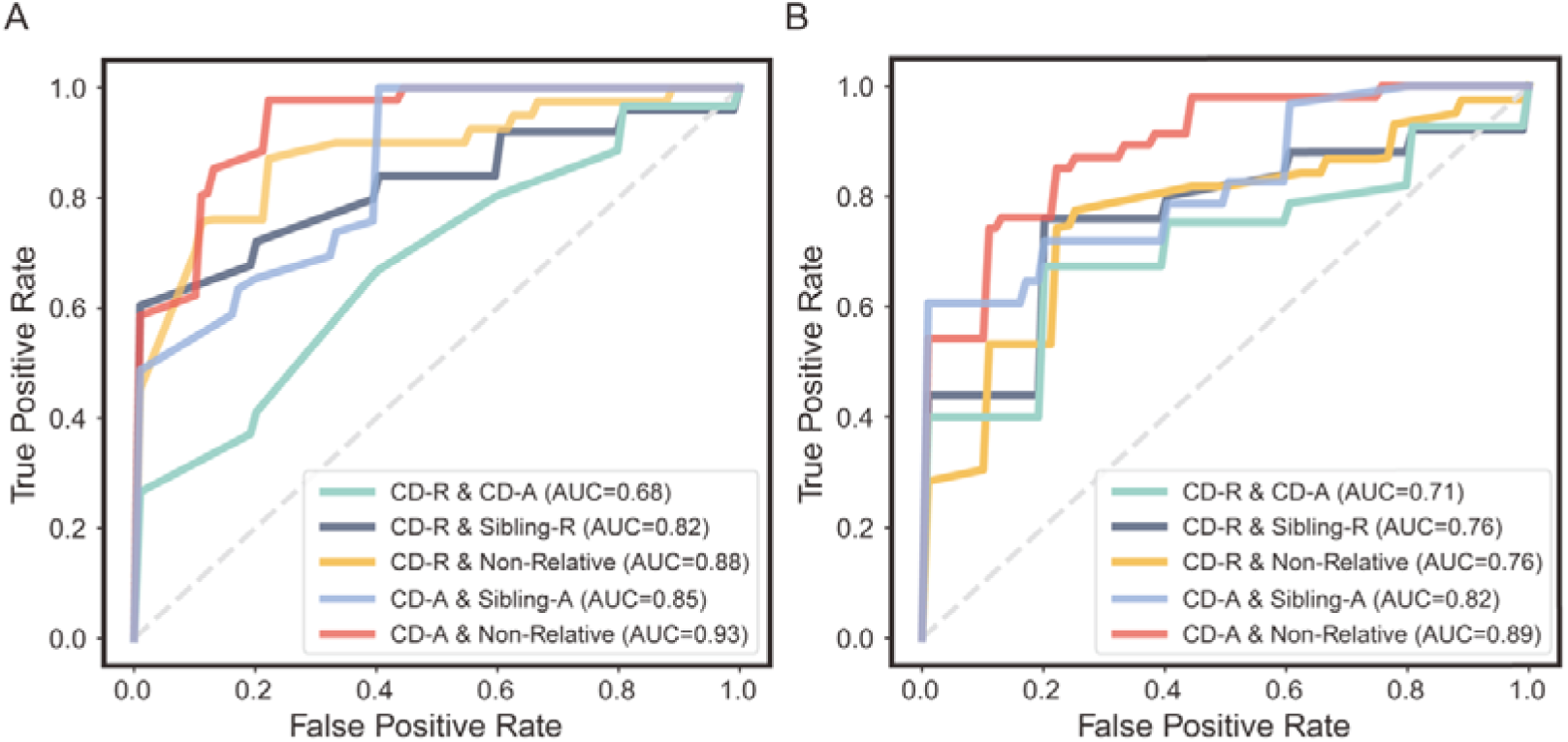
Diagnostic potential of gut microbial markers for CD. **(A)** Diagnostic performance of the random forest classification model based on the CD-R specific differential genera, *Dorea*, *Faecalibacterium* and *Fusicatenibacter*. This model was trained for distinguishing: CD-R from CD-A, CD-R from Sibling-R, CD-R from non-relative, CD-A from Sibling-A, and CD-A from non-relative. **(B)** Diagnostic performance of the Support Vector Classification model based on 74 CD-R specific differential KO genes related to SCFA production. This model was trained for distinguishing CD-R from CD-A, CD-R from Sibling-R, CD-R from non-relative, CD-A from Sibling-A, and CD-A from non-relative.

Intriguingly, *Faecalibacterium, Dorea*, and *Fusicatenibacter* also demonstrated impressive performance in distinguishing CD-A from Sibling-A and non-relative controls, with AUROC values of 0.85 and 0.93, respectively (Figure 5). However, this classification model did not perform well in differentiating between CD-R and CD-A (AUROC = 0.68, Figure 5), which aligns with the similar alteration trends observed in both the CD-R and the CD-A groups for these three genera.

### Validation of microbial features in CD with a large external multi-center cohort

We utilized a large whole-genome shotgun sequencing dataset, iHMP(Lloyd-Price *et al*., 2019), to validate our findings and test the diagnostic potential of CD-R specific microbial features in a diverse population. This dataset was generated from a multi-center cohort of CD patients with diverse cultural and geographical backgrounds, including 383 CD patients (CD-A: 76; CD-R: 307) and 190 healthy controls (HC).

Several phyla displayed trends of altered abundances in CD-A and CD-R, but no statistical significance was achieved (Supplementary Figure S5A, Supplementary Table S7). At the family level, 10 differential families were identified between the CD-A group and the healthy control group (Supplementary Table S8), among which Rikenelleaceae and Oscillospiraceae displayed decreased abundances in CD-A, consistent with the observations in the discovery cohort (Supplementery Figure S5B). In CD-R, 6 differential families were observed compared to healthy controls, among them, Ruminococcaceae displayed decreased abundance in CD-R (Supplementery Figure S5B), aligning with the findings from the discovery cohort.

At the genus level (Supplementary Table S9), 16 differential genera were identified between CD-A and healthy controls. Decreased abundances of SCFA-producing *Alistipes*, *Roseburia*, and *Butyricicoccus* were observed in CD-A, consistent with the discovery cohort (Figure 6A). On the other hand, 13 differential genera were identified between CD-R and healthy controls, among them, *Faecalibacterium* and *Fusicatenibacter* displayed decreased abundances in CD-R (Figure 6A), in line with the findings from the discovery cohort. *Dorea*, which displayed decreased abundance in CD-R in the discovery cohort, exhibited a trend of decreased abundance in CD-R in the validation cohort. It is worth noting that SCFA-producing genera *Agathobaculum* and *Eubacterium* displayed decreased abundances in CD-R (Figure 6A). Similar to the discovery cohort, no gradual alteration in abundance from healthy to CD-R and then to CD-A was observed for *Faecalibacterium* (Supplementary Figure S6), supporting the argument that the abundance of *Faecalibacterium* is not associated with the status of inflammation.

**Figure 6.**
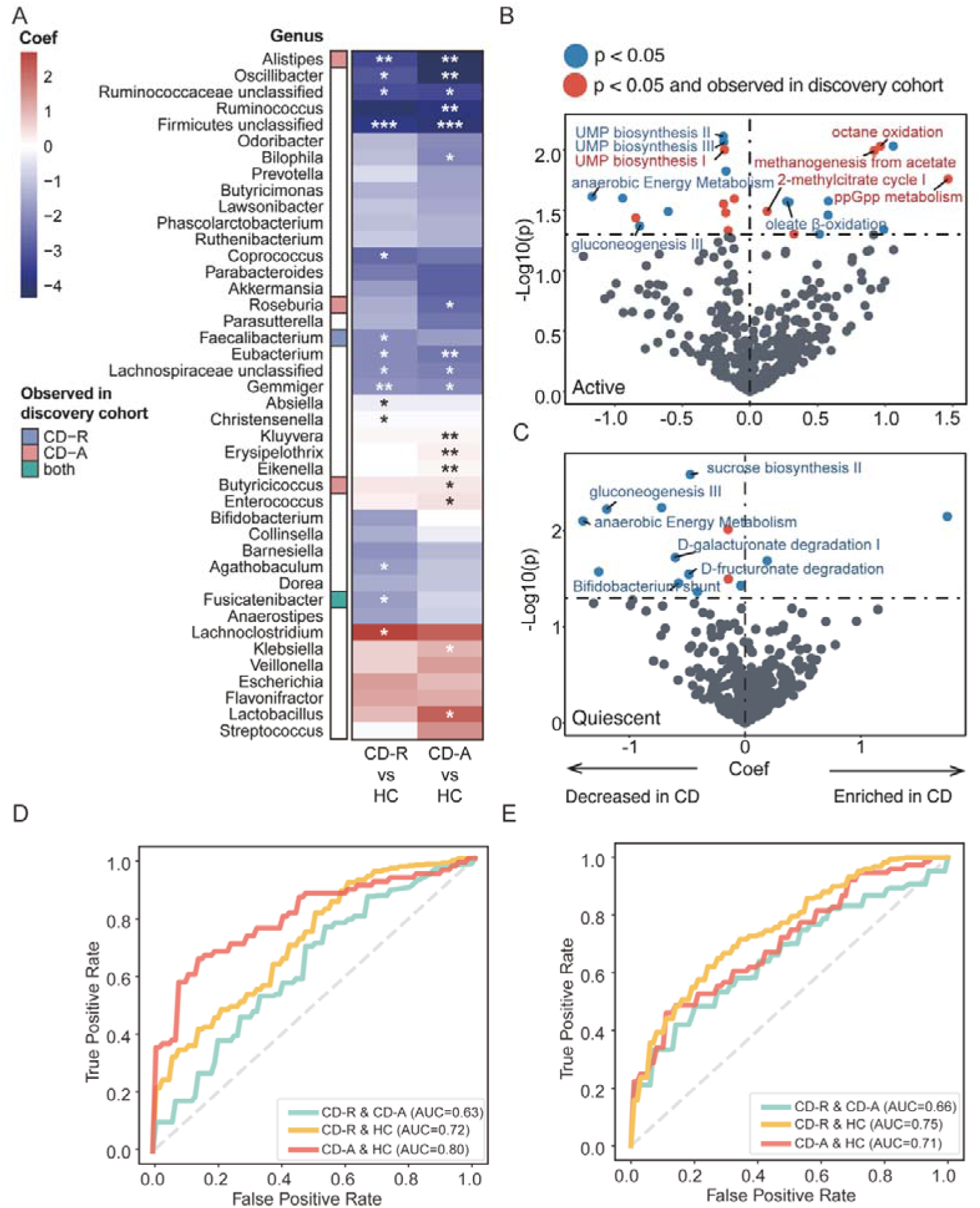
Validation of microbial features in CD with a large external multi-center cohort. **(A)** Differential genera in CD-A and CD-R compared to non-relative healthy controls (HC). The heatmap displays the effect sizes (coefficients from a linear regression model) representing the differences in relative abundance of each genus, comparing between CD-A and HC, as well as between CD-R and HC, with significance levels indicated in the heatmap blocks. *, p < 0.05; **, p < 0.01; ***, p < 0.001. Differential genera, as well as the top 25 genera based on their coefficients, are plotted. Colors in the side bar indicate whether a specified genus was identified as a differential genus in the discovery cohort. **(B)** Differential pathways in the gut microbiota between CD-A and HC. Pathways with a p < 0.05 are highlighted in the volcano plot. (C) Differential pathways in the gut microbiota between CD-R and HC. Pathways with a p < 0.05 are highlighted in the volcano plot. **(D)** Validation of the diagnostic capability of the CD-R specific differential genera. The random forest classification model based on *Dorea*, *Faecalibacterium* and *Fusicatenibacter* was tested for distinguishing CD-R from CD-A, CD-R from healthy controls, and CD-A from healthy controls, with the validation cohort. **(E)** Validation of the diagnostic capability of the CD-R specific differential KO genes. The Support Vector Classification model based on 74 CD-R specific differential KO genes were tested for distinguishing CD-R from CD-A, CD-R from healthy controls, and CD-A from healthy controls, with the validation cohort. These differential KO genes are related to SCFA production.

At the functional level, 25 pathways were differentially enriched between the CD-A and the healthy control groups (Figure 6B, Supplementary Table S10). Among them, higher representations in oxygen-requiring metabolic pathways including octane oxidation and oleate β-oxidation, while a lower representation in pathway anaerobic energy metabolism in the CD-A group reflects higher abundance of facultative anaerobes in the CD-A group, consistent with the findings from the discovery cohort. In CD-R, 14 differential pathways were identified compared to the healthy control group (Figure 6C, Supplementary Table S11). In line with the findings from the discovery cohort, pathways contributing to SCFA production including anaerobic energy metabolism and bifidobacterium shunt, displayed decreased abundances in the CD-R group.

Next, the diagnostic capabilities of *Dorea*, *Fusicatenibacter* and *Faecalibacterium* were evaluated with the validation cohort. Using a random forest classification model, the combination of these three biomarkers were able to distinguish CD-R from healthy controls, as well as CD-A from healthy controls, achieving AUROCs of 0.72 and 0.80, respectively (Figure 6D). Similar to the discovery cohort, these markers showed limited ability in distinguishing CD-A from CD-R.

Finally, in the validation cohort, we assessed the diagnostic capabilities of differential KOs between CD-R and the healthy controls. We used 74 differential KOs associated with SCFA production to develop a random forest model, which achieved an AUROC of 0.75 for distinguishing CD-R from healthy controls, and an AUROC of 0.71 in distinguishing CD-A from healthy controls (Figure 6E).

### Superior performance of the CD-R specific microbial markers identified using HFDR controls versus those identified using unrelated controls

Next, we compared the performance of the CD-R specific microbial markers we identified using HFDR controls (i.e. *Dorea, Fusicatenibacter* and *Faecalibacterium*) to those established using unrelated controls, including two microbial genera panels (marker panel 1(Braun et al., 2019) and 3(Seksik et al., 2003)) and one microbial family panel (marker panel 2(Braun *et al*., 2019)). To ensure fair comparisons, random forest classification models constructed with these marker panels were tested on eight independent cohorts. Among these cohorts, iHMP and iHMP-pilot data(Lloyd-Price *et al*., 2019; Schirmer et al., 2018) included patients with CD-A, patients with CD-R and unrelated healthy controls, so that the models were tested to distinguish between CD-A and HC, between CD-R and HC, as well as between CD-A and CD-R. Except for one case where outside marker panel 1 and our panel (“This study”) showed similar classification power with iHMP (Figure 7A), our marker panel consistently outperformed the other marker panels in distinguishing between CD-R and HC, and between CD-A and HC, with both iHMP (Figure 7A) and iHMP-pilot data (Figure 7B). The classification power of the marker panels was also evaluated using six additional CD patient cohorts where information regarding the disease status (active or remission) of CD patients was unavailable. In these cases, the models were solely tested to differentiate between CD and HC. Across all six cohorts, our marker panel consistently demonstrated satisfactory performance with AUROCs above 0.85 (Figure 7C-H). Conversely, the outside marker panels showed either inconsistent performance (marker panels 1 and 2) or poor performance (marker panel 3) (Figure 7C-H). These data demonstrate superior diagnostic potential of our CD-R specific microbial markers identified using HFDR controls compared to previously published markers identified with unrelated controls.

**Figure7.**
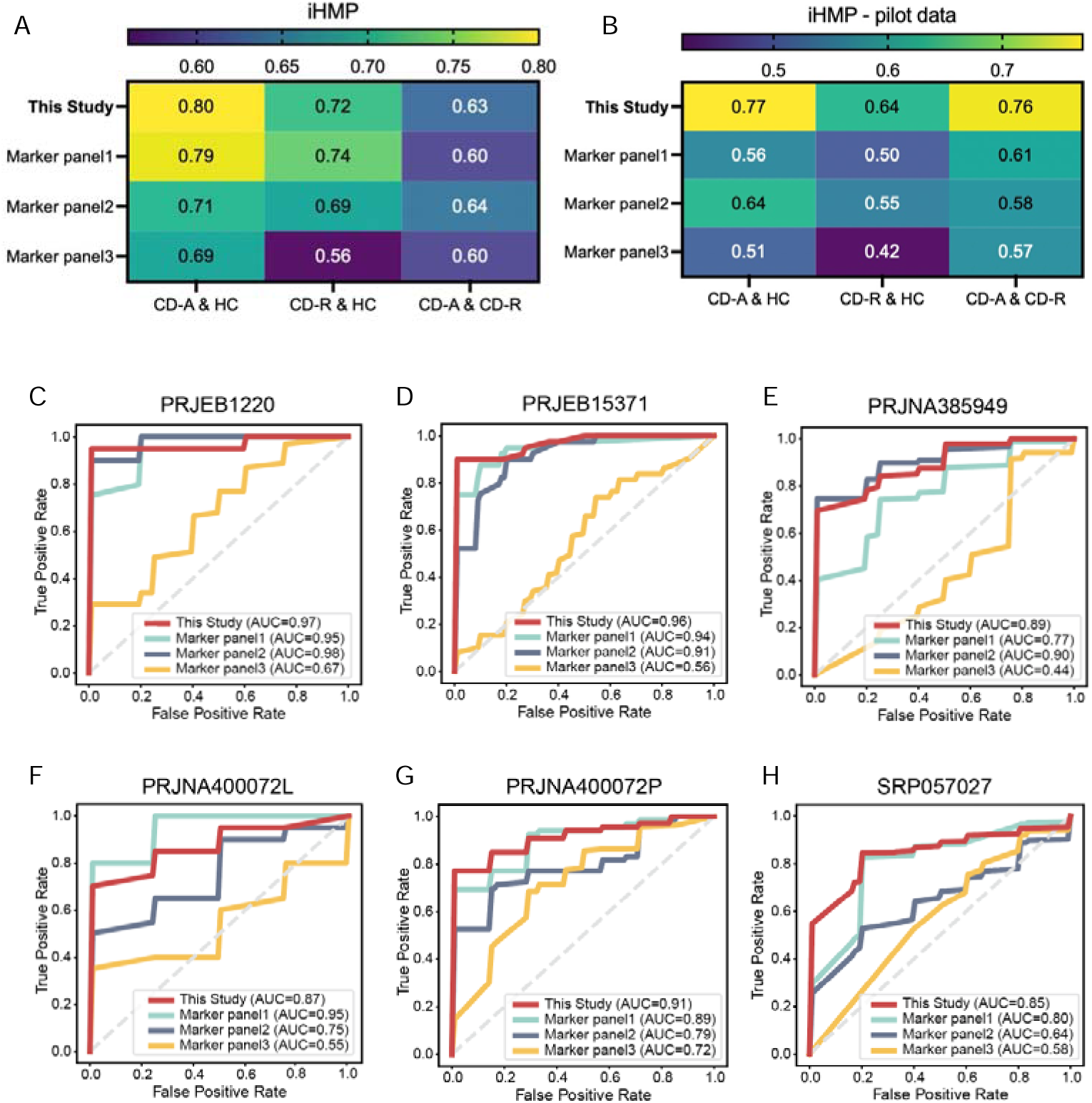
Superior performance of CD-R specific microbial markers identified using HFDRs versus those identified using unrelated controls. **(A-B)** AUROC values derived from random forest classification models, based on CD-R specific microbial markers identified using HFDR controls in this study and three other panels from published studies. These models were tested for distinguishing between CD-A and HC, CD-R and HC, as well as CD-R and CD-A, in validation cohorts iHMP (PRJNA398089) (A) and iHMP-pilot data (PRJNA389280) (B). **(C-H)** ROC curves of the random forest classification models, tested for distinguishing patients with CD from HC in additional validation cohorts, for which no information on disease status (active or remission) is available. Marker panel1, differential genera identified in Braun, T., et al.; marker panel2, differential families identified in Braun, T., et al.; marker panel3, differential genera identified in Seksik, P., et al.

## Discussion

### Using HFDRs to control for genetic and environmental confounding factors in microbiome studies of CD

In this study, we investigated changes in the fecal microbiota of patients with CD and compared them to two control groups: HFDRs and non-relative healthy individuals. Importantly, using HFDRs as controls resulted in fewer differential taxa between CD patients and controls, compared to when non-relative healthy individuals were used. This highlights a significant resemblance in gut microbiota between CD patients and their HFDRs. These findings strongly emphasize the substantial impact of genetic and environmental factors on microbial composition. Consequently, it raises an important inquiry into whether these factors play pivotal roles in CD pathogenesis or necessitate careful consideration as confounding variables.

Because the healthy relatives of CD patients are at high-risk for CD(Calkins and Mendeloff, 1986), it was proposed that the gut microbiota of healthy relatives of CD patients is a potential causal factor for the development of inflammation(Hedin et al., 2016; Jacobs et al., 2016). However, as HFDRs do not exhibit intestinal inflammation, we reason that their microbial features are not linked to, or play a role in the recurrence of intestinal inflammation. The innocence of the microbial features in HFDRs of CD patients was recently corroborated by a prospective study(Raygoza Garay *et al*., 2023). Raygoza Garay reported no significant difference in microbial taxonomic abundance between HFDRs who developed CD in five years and those remained healthy during follow-up, suggesting that the average microbiota in HFDRs does not harbor the key features driving the development of intestinal inflammation.

Considering that the dysbiosis in HFDRs is not directly linked to inflammation, it becomes crucial to use HFDRs to control for genetic and environmental confounding factors when studying the microbiota of CD patients. This strategy enables the exclusion of numerous altered taxa that are shared between CD patients and their HFDRs. As a result, it facilitates a more precise and accurate identification of specific differential taxa that are genuinely associated with intestinal inflammation.

### Evidence for the causal role of altered gut microbiota in CD-R

Using HFDRs as controls to minimize the influences of genetic and environmental confounding factors, 27 and 3 differential genera were identified in CD-A and CD-R, respectively. Several lines of evidence showed that the decreased abundances in the differential genera in CD-R, *Faecalibacterium*, *Dorea* and *Fusicatenibacter* were not consequence of inflammation, but may indeed contribute to disease relapse through decreased production of SCFAs.

Firstly, because CD-R is prone to relapses, the decreased abundance of these genera in CD-R is associated with subsequent gut inflammation, thereby indicating their potential role in CD relapse. Specifically, in both the discovery and validation cohorts, *Faecalibacterium* was not identified as a differential genus for CD-A, thereby suggesting no correlation between ongoing inflammation and *Faecalibacterium* abundance.

Secondly, while many of the differential genera showed a progressively altered abundance pattern from healthy controls to CD-R, and then to CD-A, demonstrating their association with the intestinal inflammation activities, this pattern was absent in two of the differential genera in CD-R, *Faecalibacterium* and *Dorea*. In other words, the decreased abundance of *Faecalibacterium or Dorea* in CD-R was not likely a consequence of inflammation.

Thirdly, correlations between differential genera and serum inflammation markers were prevalent in CD-A, but absent in CD-R. This further reinforces the lack of correlation between current inflammation and the differential genera in CD-R, effectively refuting the possibility that the alterations in *Faecalibacterium*, *Dorea* and *Fusicatenibacter* are a consequence of inflammation.

Lastly, fecal SCFA levels provide an additional dimension of evidence for the potential causal role of microbiota in CD-R on the relapse of intestinal inflammation. CD-R displayed trends of decreased levels in all species of SCFAs except acetate. Importantly, *Faecalibacterium*(Flint et al., 2015), *Dorea*(Taras et al., 2002) and *Fusicatenibacter*(Takada et al., 2013) are known for SCFA production, and the abundances of these genera correlated with many of the SCFAs, suggesting that decreased levels of SCFAs may underlie the pathological impact of the dysbiosis. The anti-inflammatory role of butyrate has been established through studies involving human cells and rodent colitis models(Segain et al., 2000). In line with the anti-inflammatory role of butyrate in IBD, decreased butyrate production potential has been observed in mixed populations of quiescent and active IBD cases(Frank et al., 2007; Vich Vila et al., 2018). Additionally, the decreased abundance in butyrate producing *Roseburia* is associated with higher risk scores for IBD in healthy individuals(Imhann *et al*., 2018).

Therefore, our data demonstrate associations of CD-R with decreased abundances in *Faecalibacterium*, *Dorea* and *Fusicatenibacter*, and that these alterations are not a consequence of inflammation. In addition, both microbial functional and metabolomic analyses concur in identifying decreased SCFA as a potential mechanism for disease relapse. Collectively, these observations provide multiple layers of evidence supporting a potential causal role of altered microbiota in CD-R in the relapse of intestinal inflammation. Recently, Raygoza Garay et al established a microbiome risk score (MRS) that predicts the likelihood of a healthy individual developing CD (Raygoza Garay *et al*., 2023). Their findings align with ours at the functional level, as one of the top genera contributing to the MRS is SCFA-producing *Roseburia*, and a trend of decreased abundance in *Roseburia* is linked to disease onset.

Nevertheless, the definitive causal role of the microbiota in CD remains to be determined. Further investigations, especially intervention studies, are required for establishing a conclusive relationship. In addition, future studies are required to understand the dysbiosis in CD-A, with a specific emphasis on distinguishing between altered taxa that actively contribute to sustaining inflammation and those that result from inflammation or dietary changes in CD-A.

### Potential diagnostic and therapeutic value of the microbial features in CD-R

With our internal discovery cohort and a large multi-center validation cohort with diverse geographical and cultural backgrounds, we demonstrated that microbial features in CD-R have diagnostic value for distinguishing CD from healthy individuals. Previous studies have shown the successful application of classification models based on microbial features alone(Franzosa et al., 2019; Gevers et al., 2014) or in combination with an inflammation marker(Vich Vila *et al*., 2018) to distinguish IBD patients from non-IBD controls. In this study, we took it a step further by presenting a microbiome-based classification model that achieved satisfactory performance in distinguishing both CD-R and CD-A from healthy individuals. This is particularly valuable considering the challenges in identifying CD-R using current blood and pathology tests, as well as the non-invasive nature of the microbial markers.

It is very important to note that, the CD-R specific microbial markers we identified using HFDR controls demonstrated superior performance compared to published microbial markers identified with unrelated healthy controls. Our marker panels consistently demonstrated satisfactory classification power in distinguishing between CD patients and healthy controls with multiple independent cohorts, while the performance of the outside marker panels was either inconsistent or poor. The consistent and superior performance of the markers we identified could be attributed to minimizing genetic and environmental influences during the identification of microbial markers using HFDR as controls.

To fully appreciate the significance of the performance of our diagnostic models, it is important to consider several major challenges posted by the validation cohort. Firstly, the microbial features were initially identified in the discovery cohort using 16S rRNA sequencing data with open-reference taxonomy assignment, while the validation cohort was analyzed by whole genome sequencing with closed-reference taxonomy assignment, a method leading to the exclusion of a significant number of microbes in the validation cohort. Secondly, the discovery cohort used paired HFDRs as healthy controls, whereas the validation cohort employed unrelated healthy subjects who might have been influenced by genetic and environmental factors. Lastly, the discovery cohort consisted of Chinese patients and healthy subjects, while the validation cohort comprised American and European patients and subjects. Despite these challenges, the classification model performed satisfactorily in the validation cohort, comparable to its performance in the discovery cohort. The outstanding performance observed suggests that the differential genera we identified (*Faecalibacterium*, *Dorea* and *Fusicatenibacter*) are universal features among CD patients of distinct cultural and geographical origins. Lower abundances of *Faecalibacterium* have also been observed in CD-A(Quince et al., 2015) and CD-R(Jacobs *et al*., 2016; Pascal *et al*., 2017) by other investigators. Sokol et al. reported lower representation of *Faecalibacterium* in active IBD, but not in IBD patients in remission, likely due to a small sample size(Sokol *et al*., 2009).

The differential genera used for our classification model consist of SCFA-producing bacteria. In line with this, the KO genes related to SCFA production displayed a similar diagnostic potential for distinguishing CD from healthy controls. The outstanding performance of the SCFA-related microbial features in distinguishing CD from healthy controls underscores the prevalence of diminished SCFA production capacity in the microbiota of both CD-R and CD-A, in support of their causal role in disease relapse and the perpetuation of gut inflammation. Thus, the microbial features we have identified hold promise as potential therapeutic markers. Future endeavors targeting SCFA-producing bacteria may pave the way for novel and effective therapeutics for both CD-R and CD-A.

In summary, by using HFDRs as controls to minimize the impact of genetic and environmental confounding factors, we have identified distinct microbial features in CD-A and in CD-R. Multiple lines of evidence showed that the decreased abundances in the differential genera in CD-R (*Faecalibacterium*, *Dorea* and *Fusicatenibacter*) are not a consequence of inflammation, but rather potentially play a causal role in disease relapse through decreased production of SCFAs. These microbial features within CD-R displayed outstanding potential as non-invasive markers for distinguishing CD-R and CD-A from healthy controls across diverse populations with distinct geographical and cultural backgrounds.

## Acknowledgments

The authors thank all physicians, nurses, and trainees in the Department of Gastroenterology, the Sixth Affiliated Hospital, Sun Yat-sen University for assistance in sample collection. This work was supported by the National Natural Science Foundation of China (82170542 to RZ, 92251307 to RZ, and 82270544 to MZ), the National Key Research and Development Program of China (2021YFF0703700/2021YFF0703702 to RZ), Guangdong Province “Pearl River Talent Plan” Innovation and Entrepreneurship Team Project (2019ZT08Y464 to LZ), the Sun Yat-sen University Clinical Research 5010 Program (2014008), the program of Guangdong Provincial Clinical Research Center for Digestive Diseases (2020B1111170004), and the National Key Clinical Discipline.

## Author contributions

R.Z., L.Z. and M.Z. designed and supervised the project. Y.L., W.W., L.Z. and M.Z. enrolled the discovery cohort, collected patients samples, performed the metabolomics analysis and collected clinical data. W.C. and S.G. collected the sequencing data and relavant meta-data for the validation cohort. W.C., S.G., D.W., N.J., L.Z. and R.Z. analyzed the data. W.C. and L.Z. wrote the first draft. All other authors critically revised the manuscript. All authors reviewed and approved the manuscript before submission.

## Methods

### Participants of the discovery cohort

For discovery cohort, patients with Crohn’s disease (CD), their paired healthy first-degree relatives (HFDRs), and non-relative healthy controls were enrolled as part of our study titled “<u>D</u>ietary <u>a</u>nd <u>m</u>icrobial impact <u>in I</u>nflammatory <u>B</u>owel <u>D</u>iseases with healthy first-degree relative controls (DamnIBD)”. The DamnIBD study, a single-center prospective investigation, took place at the Sixth Affiliated Hospital of Sun Yat-sen University in Guangzhou, China, between March 2014 and December 2019.

Diagnosis was were established using standard clinical, endoscopic, and histological criteria. Exclusion criteria included: any prior history of digestive tract-related diseases or surgeries other than IBD, such as gastrointestinal polyp, intestinal adenoma, and gastrointestinal tumors. Additionally, individuals who had taken antibiotics or proton pump inhibitors within one month prior to tissue collection were excluded, as well as those who lacked a healthy sibling of the patient for enrollment. Informed consent was obtained from all participants before sample collection.

Furthermore, CD patients were categorized into two subgroups based on endoscopy: CD patients with active inflammation (CD-A) and CD patients in remission (CD-R). On the day of endoscopy, blood samples were collected for blood biochemical analysis to evaluate markers for inflammation and other factors. Correspondingly, the Sibling-A and Sibling-R groups consists of HFDRs of the CD-A and CD-R, respectively. In addition, we enrolled a non-relative control group comprising healthy volunteers not related to the patients with CD. This study was approved by the Institutional Review Board of the Sixth Affiliated Hospital, Sun Yat-sen University, and informed consent was obtained from all participating subjects.

### Participants of the validation cohort

The metagenomic raw sequencing data for the iHMP cohort (CD-A: 76; CD-R: 307; HC: 190, PRJNA398089 (Lloyd-Price, et al. 2019)), iHMP pilot cohort (CD-A: 33; CD-R: 69; HC: 64, PRJNA389280 (Schirmer *et al*., 2018)), LewisJD 2015 cohort (CD: 340; HC: 26, SRP057027(Lewis et al., 2015)), FranzosaEA 2019 cohorts (CD: 20; HC: 20 for PRJNA400072L, and CD: 63; HC: 34 for PRJNA400072P (Franzosa *et al*., 2019)), HeQ 2017 cohort (CD: 53; HC: 40, PRJEB15371(He et al., 2017)), HallAB 2017 cohort (CD: 89; HC: 21, PRJNA385949 (Hall et al., 2017)) and NielsenHB 2014 cohort (CD: 23; HC: 21, PRJEB1220 (Nielsen et al., 2014)) were downloaded from the European Nucleotide Archive.

Among these validation cohorts, CD patients from iHMP cohort and iHMP pilot cohort were further stratified into CD-A and CD-R based on the patient-reported Harvey–Bradshaw index (HBI). Individuals with an HBI score ≥ 5 were assigned to the CD-A group, while those with scores < 5 were assigned to the CD-R group. Additionally, a healthy control group consisting non-IBD subjects was included. The information regarding the disease status (active or remission) of CD patients in six other CD patient cohorts was unavailable.

### Microbiota 16S amplicon sequencing and data processing

Fecal samples were collected in sterile containers and promptly stored at −80°C. Genomic DNA was extracted from these fecal samples using a stool DNA Kit (OMEGA; cat. #D4015-01), following the manufacturer’s instructions. The total DNA was stored at −80°C until its subsequent use. DNA sequencing was conducted at BGI (Shenzhen, China) using the Illumina MiSeq Benchtop Sequencer. The sequencing approach specifically targeted the V5-V6 region of the 16S rRNA gene, using a paired-end sequencing methodology.

The sequences were processed and annotated using the Quantitative Insights Into Microbial Ecology 2 (QIIME2 V.2021.04) platform(Bolyen et al., 2019). Firstly, DADA2(Callahan et al., 2016) was used to filter out low-quality sequencing reads (QL<L30). The high-quality reads were then denoised and clustered into amplicon sequence variants (ASVs) with a 100% exact sequence match. Secondly, the taxonomic assignments of ASVs were determined using a Naïve Bayes classifier(Bokulich et al., 2018) trained on sequences from the Silva-138-99 reference database. Subsequently, the representative sequences of each ASV were aligned using Fast Fourier Transform in Multiple Alignment (MAFFT)(Katoh and Standley, 2013) within the q2-phylogeny plugin. A phylogenetic tree was constructed using the Fast-Tree plugin(Price et al., 2010). Alpha and beta diversities were computed based on a rarefied count table, with a read depth set at 6,492 reads. Functional predictions, including KO genes and MetaCyc pathways, for the 16S rDNA data were carried out using Phylogenetic Investigation of Communities by Reconstruction of Unobserved States 2 (PICRUSt2)(Douglas et al., 2020).

### Metagenomic data processing

The KneadData v.0.6 tool (http://huttenhower.sph.harvard.edu/kneaddata) was employed to filter the raw data, to isolate high-quality microbial reads while eliminating contaminants. Specifically, Trimmomatic (v0.39) wrapped in kneaddata implementing parameters such as (SLIDINGWINDOW:4:20 MINLEN:50 LEADING:3 TRAILING:3) to discard low-quality reads. Furthermore, any reads aligning with the mammalian genome, bacterial plasmids, UNiVec sequences, or chimeric sequences were subsequently removed by Bowtie2(Langmead and Salzberg, 2012) wrapped in kneaddata.

Taxonomic profiles of shotgun metagenomes were generated using MetaPhlAn3(Beghini et al., 2021). This process involve the utilization of a library of clade-specific marker genes, thereby offering comprehensive microbial profiling (http://huttenhower.sph.harvard.edu/metaphlan3).

For functional profiling, we employed HUMAnN3 v3.7 (http://huttenhower.sph.harvard.edu/humann2).

### Metabolomic analysis targeting short-chain fatty acids

Fecal samples from CD patients and their paired HFDRs were subjected to metabolomics analysis targeting short-chain fatty acids (SCFA) at BGI (Shenzhen, China). Briefly, fecal samples were homogenized in phosphate buffered saline, before extraction of the SCFAs using a liquid-liquid extraction method. SCFAs were then measured using gas chromatography–mass spectrometry (GC-MS). The quantities of each SCFAs in fecal samples were measured using standard calibration curves. The method was validated by evaluating the correlation of the linearity (R^2^ > 0.99), accuracy and repeatability.

### Dietary assessment

Dietary intake for the study subjects were assessed using a 24-h dietary recall, which was obtained through face-to-face interviews. The daily intake of each macronutrient and micronutrient was then analysed using the Automated Self-Administered 24-Hour Dietary Assessment Tool (ASA24) version (2023), developed by the National Cancer Institute, Bethesda, MD (https://asa24.nci.nih.gov)(Subar et al., 2007).

### Identificatiion of altered gut microbiota and function

In the discovery cohort, we conducted the Wilcoxon signed-rank test between patients and non-relative healthy controls, as well as the paired Wilcoxon signed-rank test between patients and their HFDRs. The p-value were adjusted using the Benjamini–Hochberg method. A threshold of FDR < 0.1 and FDR < 0.05 was employed to identify differential taxa and functions, respectively. Within the validation cohort, abundances were fitted with the following linear mixed-effects model:

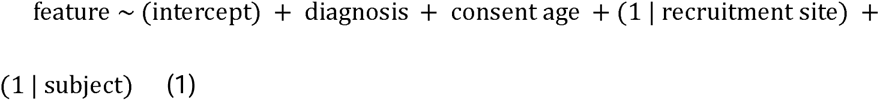

In order to mitigate the potential influence of confounding factors presented in the cohorts with non-relative controls, a linear mixed-effects model was employed. Specifically, recruitment sites and subjects were introduced as random effects to account for the correlations in repeated measures. The abundance of each feature was then characterized as a function within each phenotype (CD-A, CD-R and healthy control, with healthy control as the reference group), while adjusting for age as a continuous covariate. The coefficients derived from the aforementioned linear mixed-effects models represent the difference between the specified category and the reference category.

The fitting process was executed using MaAsLin2 (Microbiome Multivariable Associations with Linear Models 2) package in R(Mallick et al., 2021). This package employs generalized linear and mixed models to associate human health outcomes with microbial community measurements, considering the presence or absence of covariates and repeated measurements while accounting for their quantitative properties.

### Diagnostic model training and testing

We evaluated the potential diagnostic value of the CD-R specific microbial markers in distinguishing patients with different disease status. Based on the random forest (RF) model, CD-R specific microbial markers were used to construct a classification model with stratified five-fold cross-validation. Subsequently, hyperparameters such as the number of estimated trees, the maximum depth of the trees, and the numbers of features per tree of RF classifier were tuned via bayesian optimization method to optimize this classification model. And the best-performing model was built based on the final optimal hyperparameters with the highest AUC. Furthermore, we evaluated the performance of these markers against three other marker panels established using unrelated controls, including two microbial genera panels (Braun *et al*., 2019; Seksik *et al*., 2003) and one microbial family panel (Braun *et al*., 2019). This comparison was conducted across eight independent whole metagenome sequencing cohorts, utilizing the same method. Additionally, a Supporter Vector Classification (SVC) model was built using CD-R specific KO gene markers related to SCFA production. The model training and validation were executed using xMarkerFinder(Gao et al., 2022) an integrated workflow designed for microbial biomarker identification with comprehensive validations.

### Quantification and statistical analysis

#### Statistical analysis

Rare taxa present in only two or fewer samples were excluded. Subequently, the count table was transformed into relative abundance tables for further analysis. To explore associations in microbiota composition among different subjects, which encompassed patients, their corresponding healthy first-degree relatives (HFDRs), and unrelated healthy control subjects, we conducted principal coordinate analysis (PCoA) and PERMANOVA based on Bray Curties distance and unweighted Unifrac distance. Spearman correlation analysis was employed to assess relationships between serum markers, SCFA, and the differential taxa utilizing the Hmisc R package(Harrell Jr. and C., 2023).

## Data and code availability

16S rDNA sequencing data for this project are available at the Bio-Med Big Data Center (https://www.biosino.org/node-cas/, project ID: OEP004514).

Metabolomic table for this project are available at the Bio-Med Big Data Center (https://www.biosino.org/node-cas/, project ID: OEP004568)

All processed data of validation datasets for this work are available at the Bio-Med Big Data Center (https://www.biosino.org/node-cas/, project ID: OEZ014453)

Source code is available at: https://github.com/tjcadd2020/quiescent-CD-HFDR

Dietary and metabolomics data reported in this paper and any additional information required to reanalyze the data reported in this paper is available from the lead contact upon request.

Other metagenomic sequencing data used in this manuscript are available from SRA with study IDs: PRJNA398089, PRJNA389280, PRJEB1220, PRJEB15371, PRJNA385949, PRJNA400072 and SRP057027.

