## Supplementary material for "Enhanced microbiota profiling in patients with quiescent Crohn’s disease through comparison with paired healthy first-degree relatives using fecal metagenomics and metabolomics": Supplementary Figure S1.pdf

### Chen et al. Figure S1

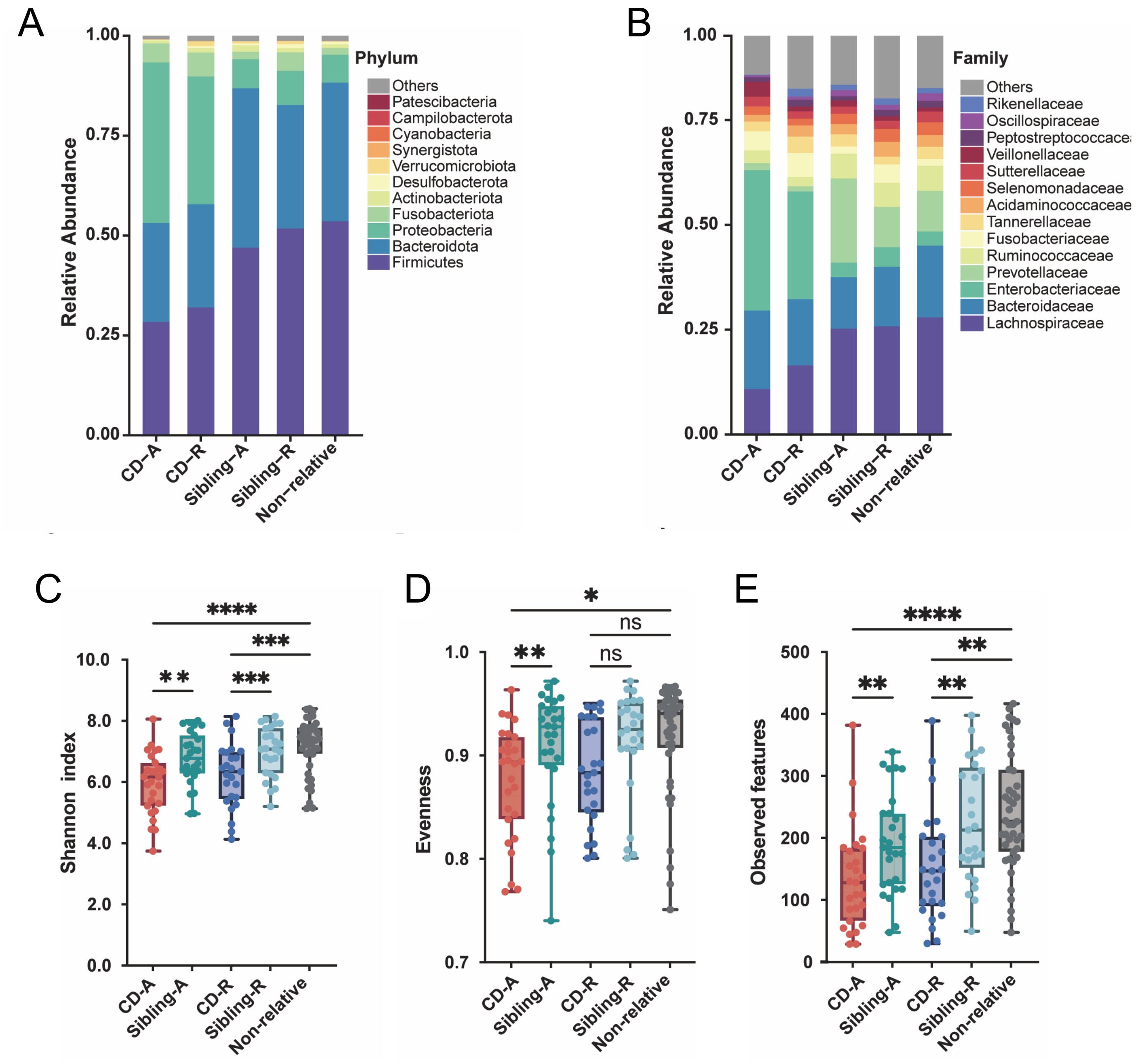


**Supplementary Figure S1 |** Gut microbial compositions and ecological diversities in patients with Crohn's disease (CD) and healthy controls.

(A) Phylum distribution of the gut microbiota in patients with active and quiescent CD (CD-A and CD-R), the healthy siblings of the patients (Sibling-A and Sibling-R), and non-relative healthy controls (Non-relative). (B) Family distribution of the gut microbiota in CD-A, CD-R, Sibling-A, Sibling-R and Non-relative groups. (C-E) Alpha diversities of the study groups as measured by Shannon index (C), species evenness (D) and Observed species (E). Paired Wilcoxon rank sum tests were performed for CD-A vs. Sibling-A and CD-R vs. Sibling-R, and unpaired Wilcoxon rank sum tests for CD-A vs. non-relative and CD-R vs. non-relative. P values were adjusted for multiple tests. *, FDR < 0.05; **, FDR < 0.01; ***, FDR < 0.001; ****: FDR < 0.0001.
