## Supplementary material for "Enhanced microbiota profiling in patients with quiescent Crohn’s disease through comparison with paired healthy first-degree relatives using fecal metagenomics and metabolomics": Supplementary Figure S2.pdf

Chen et al. Figure S2

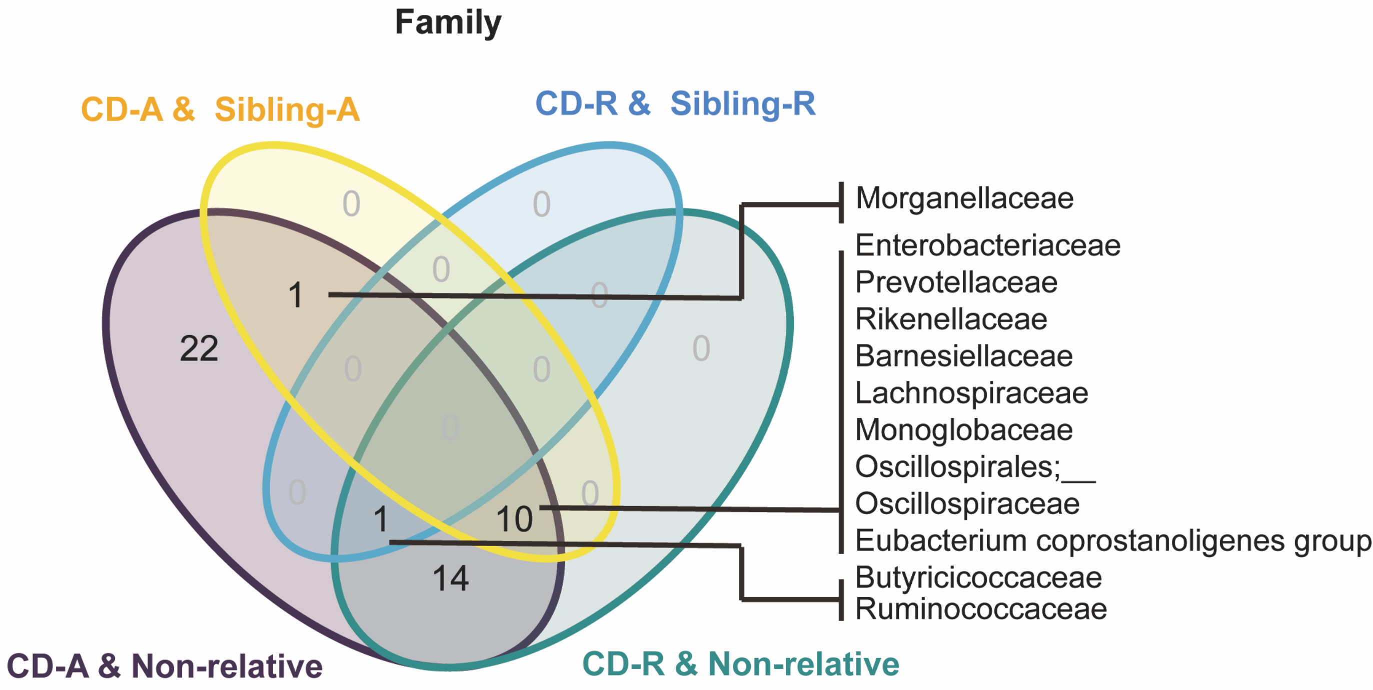


**Supplementary Figure S2 |** Venn diagram of the differential families. Included are differential families between patients with active Crohn's disease (CD-A) and paired healthy siblings (Sibling-A), patients with quiescent CD (CD-R) and paired healthy siblings (Sibling-R), CD-A and non-relative healthy controls (Non-relative), as well as CD-R and Non-relative. Names of the differential families between CD-A and Sibling-A, as well as CD-R and Sibling-R, are indicated.
