## Supplementary material for "Enhanced microbiota profiling in patients with quiescent Crohn’s disease through comparison with paired healthy first-degree relatives using fecal metagenomics and metabolomics": Supplementary Figure S3.pdf

Chen et al. Figure S3

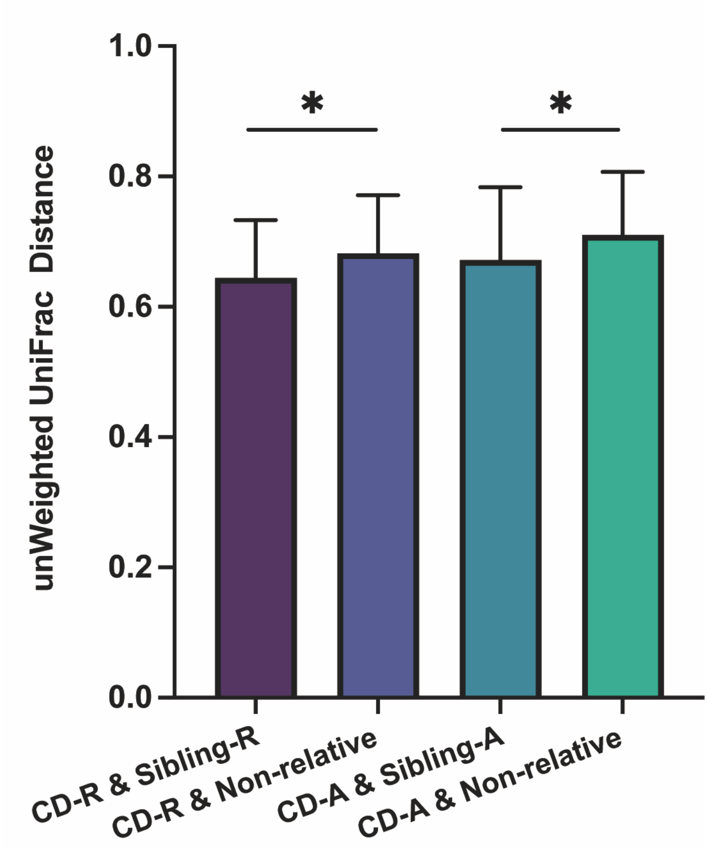


**Supplementary Figure S3 |** Unweighted UniFrac distances between study groups. Comparisons were made between patients with quiescent Crohn's disease (CD-R) and paired healthy siblings (Sibling-R), CD-R and non-relative healthy controls (Non-relative), patients with active CD (CD-A) and paired healthy siblings (Sibling-A), CD-A and Non-relative. Values are mean ± SD. *, $p < 0.05$.
