## Supplementary material for "Enhanced microbiota profiling in patients with quiescent Crohn’s disease through comparison with paired healthy first-degree relatives using fecal metagenomics and metabolomics": Supplementary Figure S4.pdf

### Chen et al. Figure S4

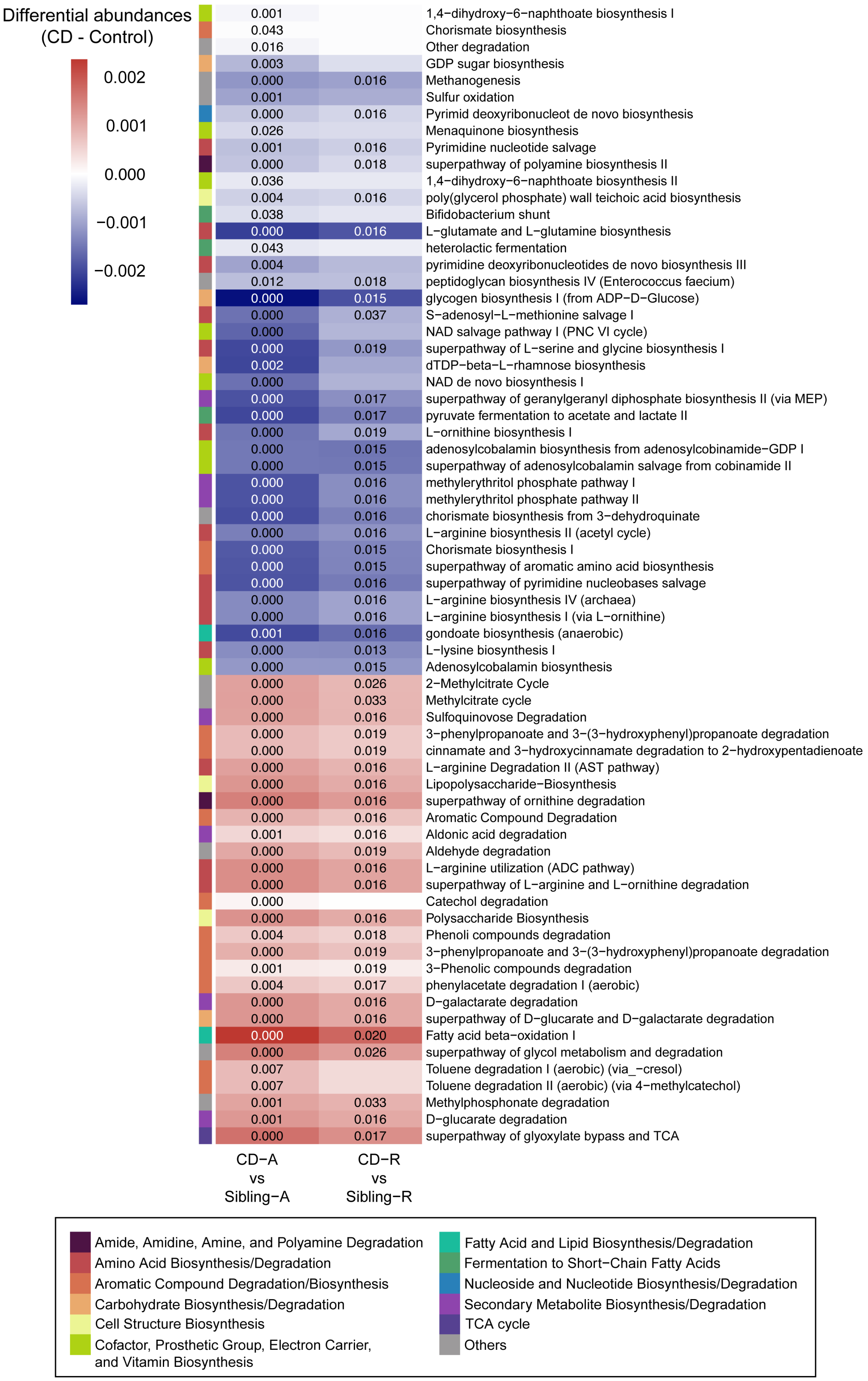


**Supplementary Figure S4 | Differential microbial pathways in the CD-A and the CD-R groups**

Differential pathways in CD-A and CD-R were identified in comparison to Sibling-A and Sibling-R, respectively. Plotted are the differential abundances (DA) of microbial pathways between CD patients and controls (“CD – control”, including “CD-A – Sibling-A” and “CD-R – Sibling-R”). The top 25 pathways with increased and decreased abundances in CD-A and CD-R are included in the heatmap. Significance levels (FDR values) are indicated in the heatmap blocks.
