## Supplementary material for "Enhanced microbiota profiling in patients with quiescent Crohn’s disease through comparison with paired healthy first-degree relatives using fecal metagenomics and metabolomics": Supplementary Figure S5.pdf

### Chen et al. Figure S5

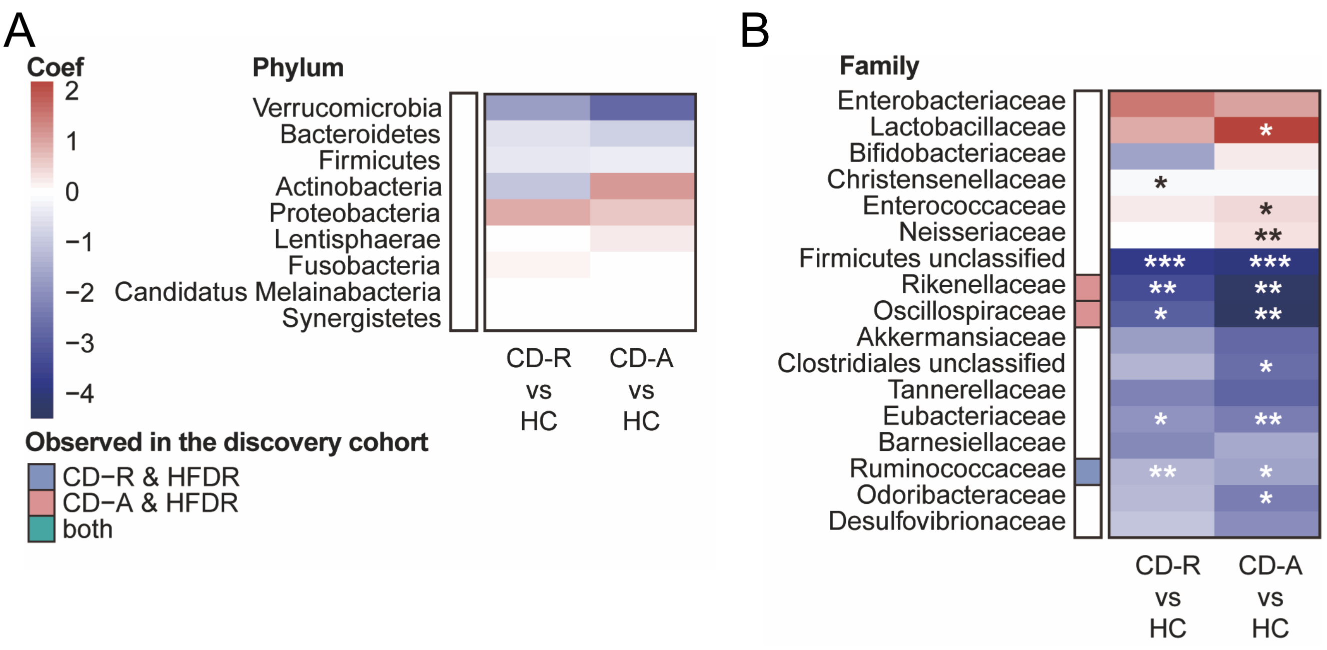


**Supplementary Figure S5 |** Compositional alterations in active and quiescent Crohn's disease in the validation cohort. (A) Differential abundances of each phylum between quiescent Crohn's disease (CD-R) and healthy controls (HC), as well as between active CD (CD-A) and HC. No differential phylum was observed in CD-R or CD-A. (B) Differential families in CD-A and CD-R compared to non-relative healthy controls (HC). The heatmap displays the effect sizes (coefficients from a linear regression model) representing the differences in relative abundance of each family. Comparisons are made between CD-A and HC, as well as between CD-R and HC, with significance levels indicated in the heatmap blocks. *, $p < 0.05$; **, $p < 0.01$; ***, $p < 0.001$. Differential families, as well as the top 15 families based on their coefficients, are plotted. Colors in the side bar indicate whether a specified family was identified as a differential family in the discovery cohort.
