## Supplementary material for "Enhanced microbiota profiling in patients with quiescent Crohn’s disease through comparison with paired healthy first-degree relatives using fecal metagenomics and metabolomics": Supplementary Figure S6.pdf

### Chen et al. Figure S6

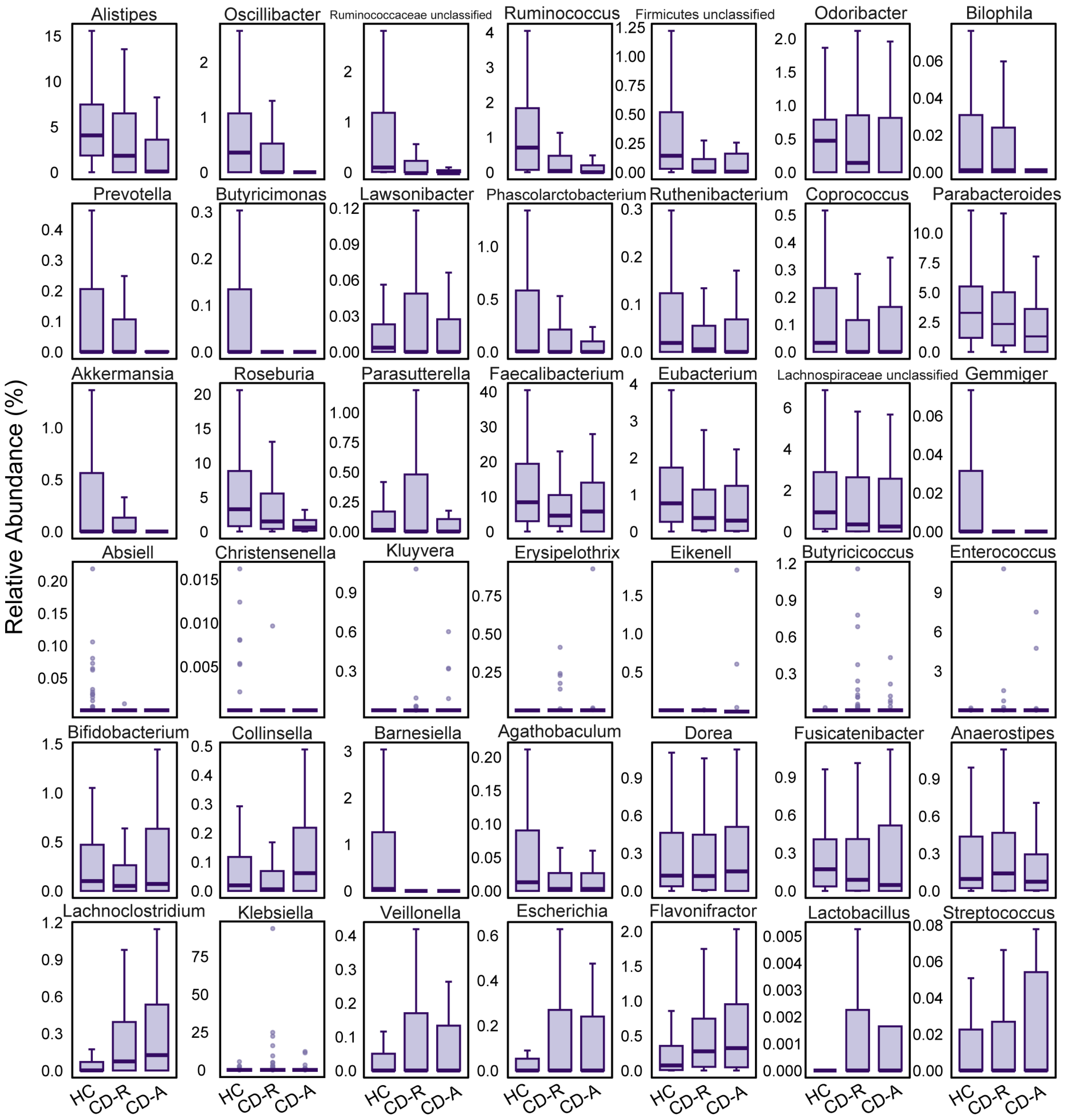


**Supplementary Figure S6 |** Relative abundances of selected genera in patients with Crohn's disease and healthy controls in the validation cohort. Included in the boxplots are differential genera, as well as the top 25 genera with the largest differences. CD-A, patients with active Crohn's disease; CD-R, patients with quiescent CD; HC, healthy controls.
